# Validating genome-wide CRISPR-Cas9 function in the non-conventional yeast *Yarrowia lipolytica*

**DOI:** 10.1101/358630

**Authors:** Cory Schwartz, Jan-Fang Cheng, Robert Evans, Christopher A. Schwartz, James M. Wagner, Scott Anglin, Adam Beitz, Weihua Pan, Stefano Lonardi, Mark Blenner, Hal S. Alper, Yasuo Yoshikuni, Ian Wheeldon

## Abstract

Genome-wide mutational screens are central to understanding the genetic underpinnings of evolved and engineered phenotypes. The widespread adoption of CRISPR-Cas9 genome editing has enabled such screens in many organisms, but identifying functional sgRNAs still remains a challenge. To address this limitation, we developed a methodology to quantify the cutting efficiency of each sgRNA in a genome-scale library in the biotechnologically important yeast *Yarrowia lipolytica*. Screening in the presence and absence of native DNA repair enabled high-throughput quantification of sgRNA function leading to the identification of high efficiency sgRNAs that cover 94% of genes. Library validation enhanced the classification of essential genes by identifying inactive guides that create false negatives and mask the effects of successful disruptions. Quantification of guide effectiveness also creates a dataset from which functional determinants of CRISPR-Cas9 can be identified. Finally, application of the library identified mutations that led to high lipid accumulation and eliminated pseudohyphal morphology.

## Introduction

A critical challenge in CRISPR-based library screens is the inability to separate active from inactive guides to ensure and quantify genome-wide coverage. Typically, multiple guide RNAs are designed to target each gene of interest with the hypothesis that some guides may not be functional ^1, 2^. This strategy maximizes the likelihood of full genome coverage through redundant targeting of each gene and has been successful in identifying new phenotypes ^3–5^, but the presence of inactive guides can obscure screening results and create false negatives ^6^. While *in silico* design and activity predictions are emerging, complete training sets that correlate CRISPR endonuclease activity, guide sequence, and local genetic context are not yet available.

Herein, we validate the genome-wide function of a synthetized library useful for CRISPR-Cas9 screens in the oleaginous yeast *Yarrowia lipolytica*. We selected this non-conventional yeast because it has value as a bioprocessing host for the conversion of biomass derived sugars and industrial waste streams (*e.g*., glycerol, alkanes, and fatty acids) into value added chemicals and fuels^7–10^. Unlike the model yeast *Saccharomyces cerevisiae*, DNA repair in *Y. lipolytica* and most other eukaryotes is dominated by nonhomologous end-joining (NHEJ)^11,12^. We reasoned that the effectiveness of each guide RNA in a high coverage CRISPR library could be quantified by comparing library evolution in an NHEJ-proficient strain and an NHEJ-deficient strain. In a strain lacking NHEJ and a homologous template (*i.e.*, NHEJ-disrupted haploid *Y. lipolytica*), double strand break repair is severely limited, and the most likely outcome of a Cas9-induced DNA double strand break is cell death^13^. Using this approach, CRISPR-Cas9 activity can be coupled to cell viability thus providing a facile, quantitative metric of guide efficiency.

Validated CRISPR libraries promise to generate more accurate and robust genetic screens with a drastically reduced false negative rate compared with libraries that are naïve to the cutting efficiency of each guide. We demonstrate this by identifying essential genes and by screening for the industrially relevant phenotypes of increased lipid accumulation and elimination of pseudohyphal cell morphology. In addition, when coupled with genome structure analysis (*i.e*., nucleosome occupancy), validation experiments can provide insights into the biological effects that dictate guide RNA effectiveness and genome accessibility. Defining a list of validated guide RNAs for any given organisms is valuable in and of itself, and is also an important step in moving towards high coverage, combinatorial screens that simultaneously target multiple genes in the genome^14^.

## Results

### Genome-wide CRISPR-Cas9 library design, construction, and analysis

We designed a library of single guide RNAs (sgRNAs) to target 7,854 coding sequences in *Y. lipolytica* PO1f (Figure 1a, Supplementary Fig. 1). Unique sgRNAs were designed to target the first 300 exon base pairs in each ORF, then scored and ranked based on their predicted on-target cutting efficiency^15^. The complete library was subsequently designed to contain the 6 highest scoring sgRNAs for each ORF, along with 480 nontargeting controls. The library was synthesized and subsequently cloned into an expression vector with sgRNA expression driven by a synthetic RNA polymerase III (Pol III) promoter^16,17^. Sequencing revealed that over 97% of the designed sgRNAs were well represented in the library (Supplementary Fig. 2).

**Figure 1.**
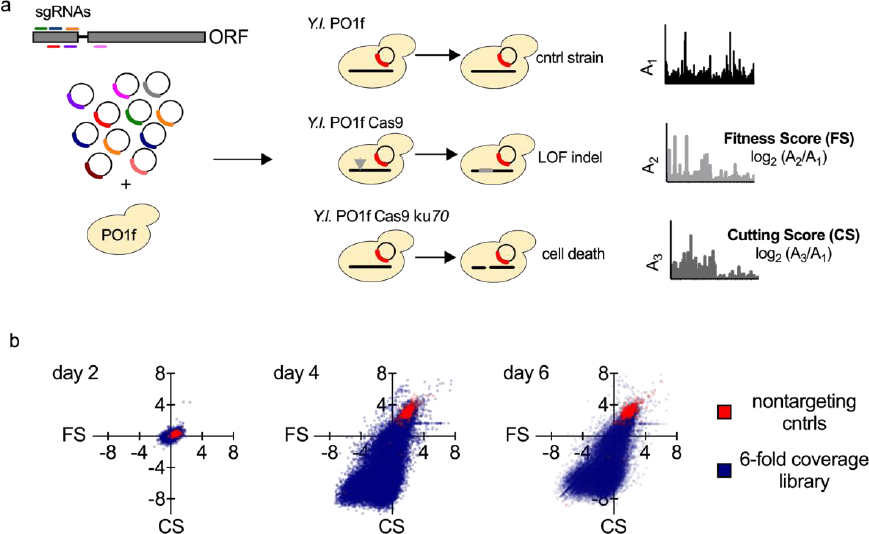
Genome-wide CRISPR-Cas9 validation and screening in *Yarrowia lipolytica*. (a) Schematic representation of sgRNA library design and workflow for genome-wide validation and screening. *Y. lipolytica* PO1f was used as the base strain for all experiments. Fitness score (FS) experiments used PO1f with constitutive expression of Cas9 from *Streptococcus pyogenes* (PO1f Cas9). Cutting score (CS) experiments used PO1f Cas9 with functionally disrupted nonhomologous end-joining (NHEJ) by inactivation of *KU70* (PO1f Cas9 *ku70*). LOF indel indicates loss of function insertion or deletion, and cntrl strain indicates the negative control strain that does not express Cas9 but does contain the sgRNA library. (b) Comparison of FS to CS for each sgRNA after 2, 4, and 6 days of growth on defined minimal media containing 2% glucose. Transformations were done in biological triplicates with at least 100-fold coverage of the library in each replicate. FS and CS values shown are the mean of the three biological replicates.

Growth screens in three different *Y. lipolytica* strains enabled us to experimentally determine two unique metrics for each sgRNA (Figure 1a). A fitness score (FS) was defined as the log2 ratio of the abundance of any given sgRNA in a Cas9 expressing strain (PO1f Cas9) to the abundance of the same sgRNA in the negative control (PO1f). Each sgRNA was also given a cutting score (CS) by calculating the same log2 ratio, but in this case comparing a Cas9 expressing NHEJ deficient strain (PO1f Cas9 *ku70*) to the control strain. FS is a measure of how harmful a CRISPR-Cas9 mediated indel was to cell growth, whereas CS is a measure of how efficient an sgRNA was for inducing a Cas9-mediated DNA double strand break. A pilot scale experiment with a nontargeting sgRNA and known functional sgRNAs targeting an essential gene and a nonessential gene confirmed that the outcomes of the FS and CS experiments were consistent with this scoring methodology (Supplementary Fig. 3).

The evolution of the full genome-wide library over three subculture cycles (6 days in total) and corresponding CS and FS values are shown in Figure 1b. Only after two growth cycles (day 4) were significant changes in the library distribution observed. The nontargeting control population behaved as expected, shifting to the upper right quadrant of the FS/CS plot indicating nonfunctional sgRNAs that resulted in no change in cell fitness. The day 4 and day 6 data indicate that the majority of the library was able to create double strand breaks. This is apparent from the shift toward the lower two quadrants. The lower left quadrant is indicative of sgRNAs that both cut (negative CS) and have a negative effect on growth (negative FS). The lower right quadrant also contains sgRNAs that cut effectively but have a neutral or positive effect on cell growth (positive FS). The upper left quadrant is only sparsely populated around the origin, likely because this quadrant represents instances with observed negative growth effects in the absence of a functional sgRNA (positive CS). Overall, three important trends are apparent: 1) the library skews strongly towards negative CS values, indicating that many of the sgRNAs are functional; 2) sgRNAs that do not produce a cut in the genome can be identified in the upper right and left quadrants; and, 3) a broad range of FS values results from outgrowth-based screens, suggesting that the library can be used to select for a wide variety of phenotypes.

### sgRNA validation and analysis of functional determinants

Based on the average CS of the nontargeting controls, we set a value of 2 as the threshold to identify inactive sgRNAs (*i.e*. noncutters). At the other end of the spectrum, we set a CS value of −5 (a 32-fold reduction in sgRNA abundance) as the lower limit of sgRNAs considered to be excellent cutters. This threshold was set based on a subpopulation of sgRNAs whose abundance was reduced to zero after two subculture cycles (4 days). Two intermediate categories were also defined: poor (−1<CS< 2) and moderate (−5<CS<−1) cutters. Figure 2a shows the CS distribution of the full library as well as six subpopulations ranked from best sgRNA per gene (1^st^) to worst sgRNA per gene (6^th^). Each subpopulation represents 1-fold coverage of the genome. The best subpopulation contained an excellent cutter for 94.6% of the targeted genes, while 82.6% of the genes have at least two excellent cutters. Histograms of each subpopulation, and the number of excellent cutters per gene are presented (Figures 2b and c, and Supplementary Figs. 4 and 5).

**Figure 2.**
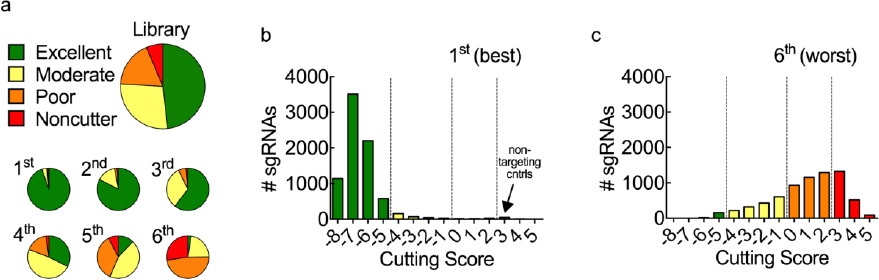
Genome coverage and distribution of CRISPR-Cas9 cutting efficiency. (a) Library and genome coverage by sgRNA classification: Excellent (CS<−5; green; 48% of full library), Moderate (−5<CS<−1; yellow; 28%), Poor (−1<CS<2; orange; 18%), and Noncutter (>2; red; 6%). The sgRNAs for each gene were ranked by CS from 1^st^ to 6^th^ and separated into six subpopulations each representing 1-fold genome coverage. (b, c) Histograms of the 1^st^ and 6^th^ sgRNA subpopulations. Mean CS values were determined after two growth cycles (4 days) from triplicate biological samples.

This CS analysis revealed that almost half of the library (48%) falls into the excellent category. Another 5% (2,828) were found to be inactive and were designated as noncutters (Figure 3a). To better understand the determinants of CRISPR-Cas9 function in *Y. lipolytica*, we looked for trends in the noncutter population. PolyT motifs (4 or more consecutive T’s) are known terminator sequences for RNA Pol III promoters, and while 1,808 sgRNA sequences in the full library contained this motif, only 492 were identified as noncutters. The CS values of these subpopulations were significantly above that of the full library (Figure 3b). The variability of CS within the polyT-containing guides suggests that while this motif can negatively impact CRISPR function, its presence does not inherently exclude the possibility of efficient cutting. sgRNAs that target near the ends of chromosomes were also found to be enriched with noncutters. The CS values of sgRNAs proximal to the end of each chromosome (731 in total; see Methods) were significantly increased relative to the library, and 463 of these were noncutters (Figure 3b and Supplementary Fig. 6).

**Figure 3.**
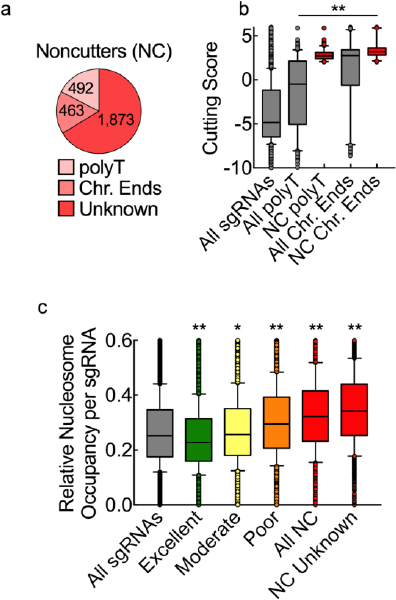
Identifying determinants of CRISPR-Cas9 function. (a) Classification and distribution of noncutting (NC) sgRNAs. (b) Cutting score (CS) of all sgRNAs (n = 46,234), sgRNAs that contain a polyT motif in the spacer sequence (All polyT; n = 1,808), noncutter sgRNAs with a polyT sequence (NC polyT; n = 492), sgRNAs proximal to chromosomal ends (All Chr. Ends; n = 731), and noncutter sgRNAs near the end of each chromosome (NC Chr. Ends; n = 463). Statistical significance was determined by oneway ANOVA with Dunnett’s multiple comparison (F-value = 1,588; P-values to all sgRNAs < 0.0001 for all subpopulations). (c) Average nucleosome occupancy per sgRNA for all sgRNAs (n = 46,234), excellent (n = 21,898), moderate (n = 12,592), poor (n = 7,993) and noncutter sgRNAs (n = 2,828), and noncutter sgRNAs that do not contain a polyT motif and are not proximal to chromosomal ends (NC Unknown; n = 1,873). Statistical significance was determined by one-way ANOVA with Dunnett’s multiple comparison (F-value = 544; P-values to all sgRNAs < 0.0001 for excellent, poor, and NC subpopulations, and p = 0.0041 for the moderate sgRNAs). Statistical significance is shown graphically with * p<0.01 and ** p<0.0001. Boxes extend from the 25^th^ to 75^th^ percentiles. The box line is plotted at the median. The whiskers extend from the 10^th^ to the 90^th^ percentiles.

The remaining 1,873 noncutters showed no obvious correlation with internal terminator sequence, chromosomal position, or predicted sgRNA secondary structure (Supplementary Fig. 7). To further evaluate these sgRNAs, we also mapped CS values to experimental estimates of nucleosome occupancy across the genome^18^. As shown in Figure 3c, excellent cutters had a lower occupancy than the library as a whole, while the unexplained noncutters had a significant increase in occupancy. Overall, nucleosome occupancy correlated well with CS (Supplementary Fig. 8). This data reveals that occupancy provides a stronger correlation to CRISPR-Cas9 activity (Pearson’s, r = 0.223) than the algorithm used to design the library^15^ (r = 0.0525) and other algorithms^19, 20^ (r = 0.0511; r = 0.1431; Supplementary Fig. 9).

### Activity validated CRISPR-Cas9 library improves essential gene analysis

Inactive sgRNAs in CRISPR libraries can produce false negatives in essential and nonessential gene classification^6^. This problem arises when one or more poor cutting sgRNAs mask the effect of successful disruptions on the FS average for a particular gene. The second dimension of data provided by CS experiments can eliminate this issue by focusing analysis on the validated cutters and excluding inactive sgRNAs. Figure 4a shows the rank-ordered FS calculated from the full library, which is naïve to the effectiveness of each sgRNA (the naïve library). Twelve nonessential genes and twelve genes that are known to be essential across eukaryotes are shown as two subpopulations (Figure 4 and Supplementary Table 1). The subpopulations partially overlapped and had closely related FS distributions when calculated from the naϊve library (Figures 4a and b). Three of the genes are notable; *ACT1*, *MYO1*, and *FOL2* are genes critical to eukaryotic cell viability, but were not distinguishable from nonessential genes, suggesting that the presence of poor cutting sgRNAs artificially increased their FS. Analysis with only excellent cutters (the validated library) resulted in a clear separation between the subpopulations: *ACT1*, *MYO1*, and *FOL2* clustered as expected with other essential genes, and the difference between FS of the essential and nonessential subpopulations increased significantly (Figures 4c and d).

**Figure 4.**
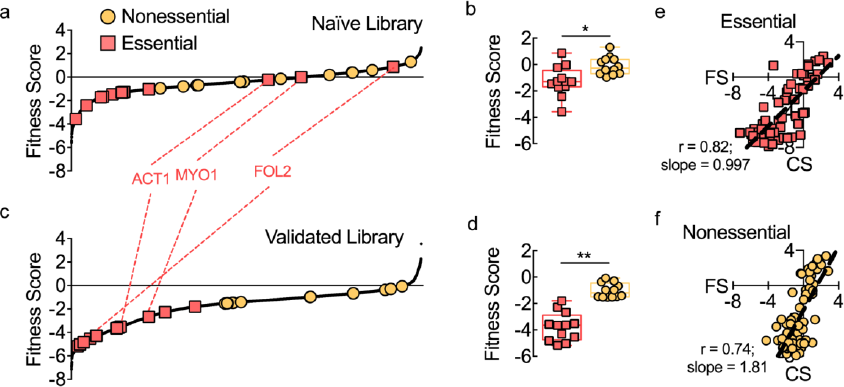
Library validation enhances essential gene identification. (a) Rank-ordered fitness score (FS) for each gene calculated using the 6-fold sgRNA library (naϊve library). Twelve nonessential (yellow) and putatively essential (red) genes are shown. (b) FS for essential and nonessential genes using the naϊve library. Boxes extend from the 25^th^ to 75^th^ percentiles, the box line indicates the median, and the whiskers extend from the 10^th^ to the 90^th^ percentiles, n = 12 for both nonessential and essential. Comparison between the means was accomplished by two-sided T-test (t = 3.029 and p = 0.0062). Statistical significance is shown graphically with * indicating p<0.01 and ** indicating p<0.0001. (c) Rank-ordered FS for each gene calculated using only the validated excellent sgRNAs (validated library). For the 5.4% of genes that did not have an excellent sgRNA, the best cutter was included. (d) FS for essential and nonessential gene subpopulations using the validated library (t = 7.764 and p < 0.0001). (e, f) Comparison of FS to CS for each sgRNA targeting the essential and nonessential subpopulations. ‘Pearson’s coefficients from linear regression (r) and slopes are shown (n = 72). FS and CS values are the mean of three biological replicates, the standard deviations associated with the means are presented in Supplementary Fig. 10.

One method of evaluating sgRNA effectiveness is to compare multiple targets to the same known essential gene^19, 21^. By definition, disruptions to essential genes are fatal (with or without intact DNA repair), which leads to a hypothesis similar to that of the CS experiments. That is, double strand breaks lead to cell death, therefore viability in a growth screen can be used as a measure of cutting efficiency. This hypothesis is supported by the observed trends in the essential and nonessential subpopulations (Figures 4e and f). The regression line of CS/FS data for all sgRNAs targeting the selected essential gene subpopulation has a slope of 0.997 (r = 0.82), suggesting a quantitative correlation between FS and CS, and providing experimental evidence to support CS as a metric for CRISPR-Cas9 activity. Appropriately, this relationship does not hold for the nonessential gene subpopulation.

The validated library identified 1,450 ORFs as essential in *Y. lipolytica*, which represents 18.4% of coding sequences (Figure 5). Using the same statistical tests, the naϊve library identified only 5.4% of the genes (424 ORFs) as essential. Genome-wide knockout collections in model yeasts suggest that ˜20% of yeast genes are essential (19.5% or 1,268 ORFs in *S. cerevisiae*^22^ and 26.1% or 1,260 ORFs in *Schizosaccharomyces pombe* ^23^), indicating that the naϊve library produced a high number of false negatives. A recent transposon-based analysis in *Y. lipolytica* supports our results^24^. The transposon screen identified 22.4% of *Y. lipolytica* genes as essential, an increase in 513 ORFs over the CRISPR-based analysis. The validated CRISPR library implicitly accounts for genomic targets that may be inaccessible due to heterochromatin structure, a known experimental limitation of transposon-based screens that can cause false positives^25^. The difference between these data sets is also likely due to challenges in transposon targeting of short ORFs (Supplementary Fig. 11).

**Figure 5.**
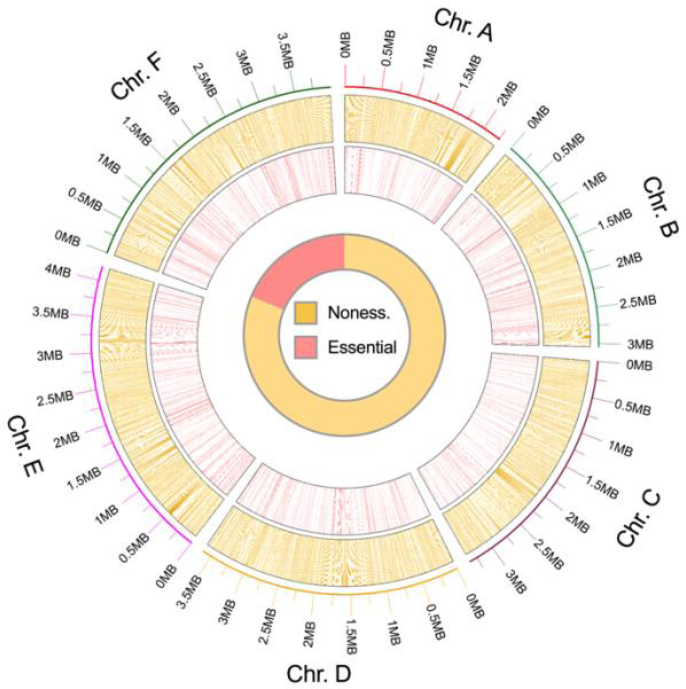
*Y. lipolytica* genome map. *Y. lipolytica* genome map with all nonessential and essential genes when grown on synthetic minimal media with 2% glucose at 30 °C are shown by chromosomal position. Nonessential genes (n = 6,404; yellow) are shown in the outer ring, and essential genes (n = 1,450; red) are shown in inner ring.

Of the 1,450 ORFs identified as essential for growth of *Y. lipolytica*, 1,003 had a homolog in *S. cerevisiae*, 502 of which were found to be essential in both organisms (Supplementary Fig. 12). 156 genes essential in *Y. lipolytica* are conditionally essential in *S. cerevisiae* for respirative growth. As *Y. lipolytica* is an obligate aerobe, respiration is essential. 98 more ORFs are duplicated in the *S. cerevisiae* genome, and 71 are involved in amino acid or nucleotide biosynthesis. GO term analysis^26^ was also performed on all genes that had a *S*. *cerevisiae* homolog. Genes involved in transcription, translation, cellular organization, and cell cycle were found to have significantly lower FS (Supplementary Fig. 13).

### Genome-wide screening for growth and nongrowth associated phenotypes

To demonstrate the utility of the CRISPR library, we performed a series of growth and nongrowth associated screens to identify genes linked to new phenotypes. The classic genetic test for canavanine resistance was successful. Ninety-four percent of the sequenced colonies that survived the canavanine challenge (50 mg/L over 2 days of growth) harbored an expression vector encoding a *CAN1* targeting sgRNA (Figure 6a). All identified sgRNAs were excellent cutters.

**Figure 6.**
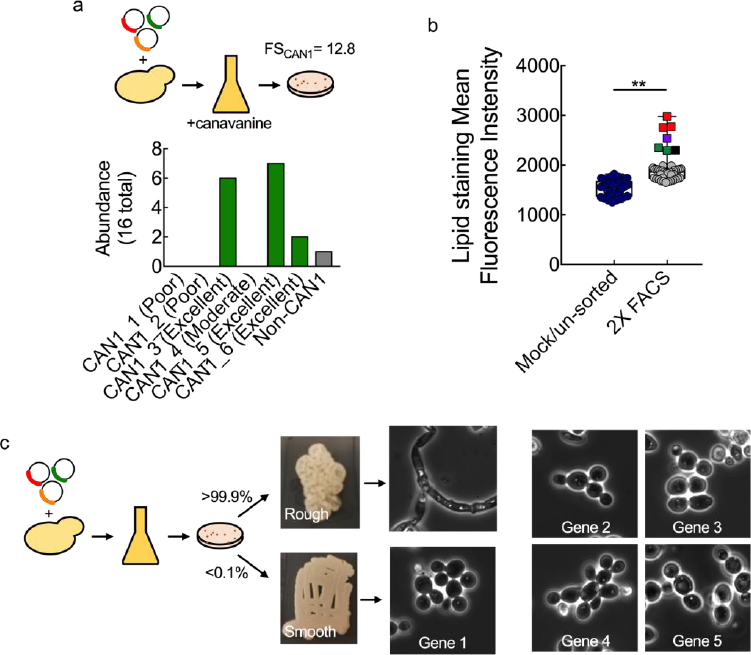
Genome-wide CRISPR-Cas9 screens for growth and non-growth associated phenotypes. (a) Growth screen for canavanine resistance. Cells were grown in the presence of 50 mg L^−1^ canavanine for 3 days, plated and sequenced. (b) Two populations were stained and analyzed using fluorescence activated cell sorting (PO1f with library and PO1f Cas9 with library). The PO1f Cas9 strain with library was sorted twice for high lipid producers to yield a high lipid producing population (2x FACS Pool). Several strains from the high lipid producing population were isolated and compared to a mock/un-sorted control consisting of cells subjected to the same produce but without a fluorescence sorting cutoff. Comparison between the means was accomplished by two-sided T-test (n = 34, 41; t = 6.713; and, p < 0.0001). Statistical significance is shown graphically with * indicating p<0.01 and ** indicating p<0.0001. (c) Non-growth associated screen for altered cell morphology. Plated colonies were visually inspected for a smooth colony appearance, indicating a lack of pseudohyphal cell morphology. 5 of ˜50,000 colonies were identified with a smooth phenotype. Rough and smooth colony appearance along with brightfield microscopy images are shown.

In a second screen, *Y. lipolytica* strains with increased lipid staining were isolated from the CRISPR library using a fluorescent lipid dye and fluorescence activated cell sorting (Figure 6b). A total of four unique genes associated with a high lipid staining phenotype were identified in this screen. Two of the four identified genes have known roles in lipid turnover and growth on fatty acids in other yeast species^27, 28^. A third gene is likely involved in the enzymatic processing of acyl-CoA species^29^, a function that is often intertwined with lipid metabolism (*e.g*., β-oxidation). The final gene in the set has no known function or similarity to a well-annotated sequence, but two different excellent sgRNAs targeting the gene were isolated from the high lipid staining subpopulation. This redundant targeting supports the validity of the result and demonstrates the utility of the screen to identify novel genotype to phenotype relationships.

A third screen tested the ability of the library to screen for a nongrowth associated phenotype. *Y. lipolytica* is a dimorphic yeast, where the transition from yeast to pseudophyphal morphology is often precipitated by environmental stress or nutrient limitations^30^. Multiday growth on solid media is often sufficient to trigger this transition, which can be readily observed by visual inspection of agar plates (Figure 6c). To identify strains deficient in pseudohyphal formation, we plated the library, identified colonies with altered morphology, confirmed cellular phenotype by microscopy, and sequenced. A total of 5 unique genes were identified as necessary for the pseudohyphal transition. These results demonstrate the utility of this library to rapidly identify novel targets for phenotypes of interest.

## Discussion

With genome-scale mutagenesis technologies such as MAGE, TRMR, Tn-seq and other similar approaches, it is now possible to generate and track genome edits and their effects on native and engineered phenotypes^31–35^. Genome-wide CRISPR systems are complementary to these approaches^3–5, 36, 37^ and can have the added functionality of genome targeted site saturation mutagenesis^38^–^41^. However, these techniques have not yet been widely translated to non-model and other industrially and medically relevant organisms. The widespread adoption of CRISPR-Cas9 promises to enable genome-wide engineering and screening for a vast array of hosts^3, 5, 12, 42–45^. Here, we designed and applied such a system for functional genomic screening and non-native phenotype engineering in the yeast *Y. lipolytica*. Key to the success of the system was our ability to functionally disable native nonhomologous DNA repair, which allowed quantification of the cutting efficiency of each sgRNA-Cas9 complex in parallel and yielded a list of validated sgRNAs for nearly every gene in the *Y. lipolytica* genome.

The additional genome-wide data created by cutting score (CS) experiments also generates new information that can be used to identify the determinants of CRISPR-Cas9 function, as well as improve screen accuracy. Our experiments in *Y. lipolytica* revealed that 48% of guides in the 6-fold coverage library were highly active, and that 94.6% of protein coding sequences were targeted by at least one excellent cutting sgRNA (Figure 2). This library validation also substantially improved essential/nonessential gene classification (Figures 4 and 5) by significantly reducing false negatives that arise in the naϊve library. Nongrowth associated screens were also successful in identifying novel gene disruptions for the industrially-relevant phenotypes of high lipid accumulation and elimination of pseudohyphal growth (Figure 6). Correlation of CS data with nucleosome occupancy and several sgRNA scoring algorithms revealed that occupancy was a stronger predictor of function than *in silico* design (Figure 3; see Refs. ^15, 18–20, 46^). We also observed that chromosomal ends were difficult to target. Current scoring algorithms have been limited by training set quality and size, relying on subsets of essential genes and/or high expression reporters to learn about the relationships between guide RNA sequence and resultant Cas9 activity^15, 19, 21^. The genome-wide activity measurements obtained in the CS experiments provide a substantially richer data set for correlating guide sequence parameters to system-level characteristics. We envision that our experiments in *Y. lipolytica* may also be possible in other eukaryotes by genetically disrupting or chemically inhibiting native nonhomologous DNA repair^47^. As such, the approach described here not only resulted in a new validated screening tool for *Y. lipolytica*, but also represents a generalizable methodology for enhancing CRISPR-based screens in other organisms.

## Author Contributions

CS and IW conceived the study. CS, CAS, and IW designed the library. JFC, RE, and YY constructed the library. CS, WP, SL, and IW performed library validation. CS, SA, AB, MB, and IW designed, completed and analyzed the essential gene experiments. CS, JW, HA, and IW completed and analyzed phenotypic screens. All authors wrote and edited the manuscript.

## Acknowledgements

This work was supported by DOE Joint Genome Institute grant CSP-503076 (IW, HA, and MB) NSF 1706545 (IW). HA and JW were supported by the Office of Naval Research (ONR) under grant N00014-15-1-2785. JW acknowledges additional support from the National Science Foundation (NSF) Graduate Research Fellowship Program (DGE-1110007). SA and MB were supported by NSF 1706134. SL and WP were supported by NSF 1526742. The work conducted by the U.S. Department of Energy Joint Genome Institute, a DOE Office of Science User Facility, is supported under Contract No. DE-AC02-05CH11231.

## Methods

### Strains and media

All strains were derived from *Yarrowia lipolytica* strain PO1f (MatA, leu2-270, ura3-302, xpr2-322, axp-2)^48^. The PO1f Cas9 strain was constructed by markerless integration of a UAS1B8-TEF(136)-Cas9-CycT expression cassette into the A08 site^17, 49^. The PO1f Cas9 *ku70* strain was constructed by disrupting *KU70* using CRISPR-Cas9 as previously described^16^.

Yeast culturing was done at 30 °C, unless specifically noted otherwise. Yeast strains were grown in YPD (1% Bacto yeast extract, 2% Bacto peptone, 2% glucose) for nonselective growth. Cells transformed with sgRNA-expressing plasmids were selected in media deficient in leucine (SD-leu; 0.67% Difco yeast nitrogen base without amino acids, 0.069% CSM-leu (Sunrise Science, San Diego, CA), and 2% glucose). For canavanine resistance screening, 50 μg/mL was added to SD-leu. For lipid overproduction experiments, SDL-leu (0.67% Difco yeast nitrogen base without amino acids, 0.069% CSM-leu, and 8% glucose) was used for outgrowth.

### Plasmid construction

For integration of Cas9, pIW1009 (UAS1B8-Cas9 integration into A08 site) was constructed. pHR_A08_hrGFP (Addgene #84615) was digested with BssHII and NheI, and Cas9 was inserted via Gibson Assembly after PCR via Cr_1250 and Cr_1254 from pCRISPRyl (Addgene #70007). To facilitate cloning of the library of sgRNAs, pIW771 (SCR1’-tRNA-AvrII site) was generated by digesting pCRISPRyl with BamHI and HindIII and circularizing.

### sgRNA library design

Custom Matlab scripts were used to design each sgRNA in the library based on a recent assembly of the *Y. lipolytica* PO1f parent strain genome, W29^50^. sgRNAs were designed for all non-redundant protein coding sequences. The top 6 sgRNAs, as ranked by on target efficiency score^15^ that met the following criteria were selected for each gene: (1) uniqueness of target sequence and PAM (final 12 bp + NGG) in genome, (2) located within first 300 bp of gene, and (3) target sequence present in both genome and mRNA sequence (to avoid targeting introns). 480 sgRNAs of random sequence, confirmed to not target in the genome, were included as nontargeting controls.

### sgRNA library cloning

Four oligo pools consisting of 25% of the designed sgRNAs each were ordered from Twist Bioscience. Equal volumes of the oligo pool (1 μM) and a complementary primer (5 μM) were mixed together and annealed using the Thermo annealing advanced protocol and the calculated Tm of the complementary primers using IDT’s OligoAnalyzer 3.1. For libraries 1 and 3, second strand synthesis reactions were completed using T4 DNA polymerase (NEB) and sgRNA-Rev2, gel extracted, and purified using Zymo Research Zymo-Spin 1 columns. Libraries 2 and 4 were amplified via Q5 DNA polymerase (NEB) using 60mer_pool-F and spacer-AarI.rev or pLeu-mock-sgRNA.fwd and sgRNA-Rev2 with 0.2 picomoles of DNA as template for 7 cycles and column purified.

For libraries 1, 3, and 4, Gibson Assembly HiFi HC 1-step Master Mix (SGI-DNA) were used to clone into pIW771 digested with AarI. Library 2 was digested with AarI and cloned into pIW771 digested with AarI with a GoldenGate reaction using T4 DNA ligase (NEB). Cloned DNA was transformed into NEB 10-beta *E. coli* and plated. Sufficient electroporations were performed for each library to yield a >10-fold excess colonies for the number of library variants, and the plasmid library was isolated.

### Yeast transformation and screening

Transformation of *Y. lipolytica* with the sgRNA plasmid library was done using a previously described method, with slight modifications^16^. 2 mL of YPD was inoculated with a single colony of the strain of interest and grown in a 14 mL tube with shaking at 200 RPM for 24-30 h. 1.5 mL of cells were washed with transformation buffer (0.1 M LiAc, 10 mM Tris (pH=8.0), 1 mM EDTA) via centrifugation at 4,000 *g* and resuspended in 600 μL transformation buffer. To these cells, 18 μL of ssDNA mix (8 mg/mL Salmon Sperm DNA, 10 mM Tris (pH=8.0), 1 mM EDTA), 90 μL of β-mercaptoethanol mix (5% β-mercaptoethanol, 95% triacetin), and 4 μg of plasmid library were added, mixed via pipetting, and incubated. After incubation at room temperature for 30 min, 900 μL of PEG mix (70% w/v PEG (3,350 MW)) was added and mixed via pipetting, and incubated at room temperature for 30 min. Cells were then subjected to heat shock for 25 min at 37 °C, washed with 10 mL H2O, and used to inoculate 25 mL of SD-leu media for outgrowth experiments. Dilutions of the transformation were plated on solid SD-leu media to calculate transformation efficiency. Transformations were pooled as necessary to ensure adequate diversity to maintain library representation (100x coverage, 5 x 10^6^ total transformants per biological replicate).

Outgrowth was allowed to proceed in 25 mL of liquid media in a 250 mL baffled flask. After 2 days, cells reached confluency (OD600 > 10), and approximately 150 μL (at least 200-fold library coverage) were used to inoculate 25 mL of fresh media as desired. Also at each timepoint, 1 mL of culture was taken, DNase I (New England Biolabs) and the corresponding buffer were added, and the mix was incubated for 1 h at 30 °C to digest any plasmid DNA in the media. Pellets were then frozen and stored at −80 °C for future analysis. Each growth cycle from inoculation to sampling represented approximately 7 cell doublings.

### Library isolation and sequencing

Pellets were thawed and resuspended in 1 mL H2O. Cells were split into 5 samples of 200 μL, and plasmid was isolated using a Zymo Yeast Miniprep Kit (Zymo Research). Samples from a single pellet were pooled, and plasmid copy number was quantified using RT-qPCR with M13_F and M13_R and SsoAdvanced Universal SYBR Green Supermix (Biorad) and verified to be higher than 1 x 10^7^.

To prepare samples for next generation sequencing, isolated plasmid was subjected to PCR using forward (Cr_1665-1668) and reverse primers (Cr_1669-1673 and Cr_1709-1711) containing all necessary sequences and barcodes^1^ (Supplementary Table 2). At least 1 x 10^7^ plasmids were used as template, and PCR reactions were not allowed to go to completion to avoid biased amplification. PCR product at 250 bp was gel extracted, samples were pooled at equimolar amounts, and submitted for sequencing on a NextSeq500 at the UCR IIGB core facility.

Next generation sequencing results were demultiplexed and mapped to each sgRNA using custom scripts. A total of 453 sgRNAs were not present in the sequencing data. In addition, 440 sgRNAs had only a very low number of reads (less than 10) in the untransformed library or were erroneously designed, and were excluded from downstream analysis. Pairwise comparison between normalized read abundances for biological replicates were done to verify consistency (Supplementary Fig. 14).

### Cutting and fitness score analysis

To calculate fitness score (FS) and cutting score (CS) for each replicate, the number of reads for each sgRNA in each sequencing sample was normalized to the total number of reads, with a pseudo-count of 0.5 added when no reads were mapped to a given sgRNA for a given biological replicate. Normalized read counts for biological triplicates were then averaged together. The log2 for each abundance was then taken, and then the FS was calculated by subtracting the PO1f log2 abundance from the PO1f Cas9 log2 abundance. CS was calculated in analogous way, but with PO1f Cas9 replaced with PO1f Cas9 *ku70*.

### sgRNA analysis

Analysis of sgRNA characteristics was done using a range of publicly available tools and data. Nucleosome occupancy was determined by mapping sgRNAs to previously published nucleosome occupancy data^18^, adding the occupancy for each base pair in the target sequence, and dividing by 20. The scoring for predicting on-target activity for each sgRNA was carried out using previously published ^15, 19, 20^. Secondary structure was predicted using the default settings for RNA using Quikfold on the DINAMelt web server (http://unafold.rna.albany.edu/)^51^. polyT sequences were identified by searching each sgRNA sequence for 4 consecutive “T”. sgRNAs at chromosome ends were annotated as the 50 sgRNAs nearest either end of each chromosome, except for the end of chromosome A (84 sgRNAs), the beginning of chromosome C (126 sgRNAs), and the end of chromosome E (71 sgRNAs). Ends of chromosomes determinations were informed by where sgRNA cutting scores increased.

### Essential gene identification

Essential genes were identified based on FS. A two-sided T-test was used to determine which genes had a FS lower than the highest of the 12 putative essential genes plus two standard deviations (*POL1* (YALI_E14644), FS<−1.38 for validated library). Samples with p values < 0.05 were considered significant. A second two-sided T-test was performed to identify which genes had a FS lower than the 12 putative nonessential genes (p<0.05), with a Bonferroni correction for multiple comparisons. The same series of tests were also applied to the FS obtained from analysis with the 6-fold coverage library (the naϊve library). In this case, the initial FS cut off was from *FOL2* (YALI1_A10254, FS<1.53).

### Essential gene comparison to *S. cerevisiae*

Mapping of all *Y. lipolytica* genes to *S. cerevisiae* genes was carried out using BLAST (E<10^−30^). Essential genes in *S. cerevisiae* were those annotated as being “inviable” null mutants^52^.

### Canavanine screen

Glycerol stocks of the transformed library taken from day 2 cultures were grown in SD-leu for 2 days, and then used to inoculate SD-leu with 50 μg/mL canavanine. Cultures were then allowed to grow to confluency and plated on solid SD-leu. Single colonies were subjected to colony PCR using Cr_1742 and Cr_1743 to identify the sgRNA expressed in each colony.

### Morphology screen

Glycerol stocks of the transformed library taken from day 2 cultures were grown in SD-leu for 2 days, and then plated on solid SD-leu. Approximately 50,000 colonies were examined for a smooth morphology after 4 days of growth on solid media. Colonies were restreaked, and smooth morphology confirmed. All colonies with confirmed smooth morphology were observed using an Olympus BX51 microscope (UPlanFL 100X 1.30 oil-immersion objective lens), and images were captured with a Q-Imaging Retiga Exi CCD camera and processed using Cellsense Dimension 1.7 software.

### LipidTOX Deep Red Staining, Fluorescence Activated Cell Sorting (FACS), and Flow Cytometry Confirmation

CRISPR libraries were cultivated for FACS in media and conditions that were previously established for *Y. lipolytica* lipid characterization^10^. Specifically, cultures were started from frozen glycerol stocks (PO1f with CRISPR library and PO1f Cas9 with CRISPR library, starting from day 2 stocks) by inoculating 50 mL of SD-leu media in a 250 mL baffled shake flask, followed by 2 days of outgrowth at 28°C. The resulting inoculum cultures were then used to start 50 mL lipid production cultures in SDL-leu media (initial optical density of 0.1) with 80 g/L glucose in 250 mL shake flasks. These shake flask cultures were grown using air-permeable plugs by shaking for 3 days at 28°C and 225 RPM in a shaking incubator (New Brunswick Scientific I 26). HCS LipidTOX Deep Red (Thermo Fisher Scientific) was then used to fluorescently stain lipids according to the manufacturer-recommended protocol: cells were pelleted, all culture media was removed, and then the cell pellet was re-suspended in 100 μL of a 1:200 dilution of lipid stain in phosphate buffered saline (PBS). Two successive rounds of fluorescence activated cell sorting (FACS) were performed on a BD FACS Aria III using the APC laser/filter set. For high lipid sorting, the top 0.3% of 3×10 ^6^ APC ^+^ cells and the top 3% of 1×10 ^5^ APC ^+^ cells were collected. After the first APC sort, the collected cells were outgrown in SD-leu to saturation (3 days) and glycerol stocked. Fresh cultures were then started from the frozen glycerol stock by inoculating 50 mL of SD-leu media in a 250 mL baffled shake flask, followed by 2 days of outgrowth at 28°C. The resulting inoculum culture was then used to start a 50 mL lipid production culture in SDL-leu media with 80 g/L glucose (initial optical density of 0.1) in a 250 mL shake flask. This shake flask culture was grown using air-permeable plugs by shaking for 3 days at 28°C and 225 RPM in a shaking incubator (New Brunswick Scientific I 26) until ready for the second sort. After the second APC sort, the collected cells were outgrown in SD-leu to saturation (3 days) and glycerol stocked again. A mock-sorted / un-sorted pool was also generated in parallel throughout this double sorting process by passing the same pool of cells through the same FACS process/conditions without applying the sorting gate (i.e. all cells were collected, regardless of APC fluorescence).

From the 2X sorted (2X FACS) glycerol stock and mock/un-sorted glycerol stock, individual colonies were then isolated on SD-leu plates to isolate individual strains. 48 colonies from the 2X FACS and 48 colonies from the mock/un-sorted plates were then subjected to colony PCR and Sanger sequencing of the sgRNA cassette. Clones with sgRNAs that were successfully colony PCR amplified, sequenced, and identified as cutters (see cutting and fitness score analysis above) were subsequently re-screened to establish a lipid staining mean fluorescence intensity for individual clones (41 from 2X FACS, 34 from mock/unsorted) using the same procedure as for FACS, but with a BD LSRII Fortessa analytical flow cytometer for quantifying APC channel fluorescence.

## Supplementary Information

**Supplementary Figure 1.**
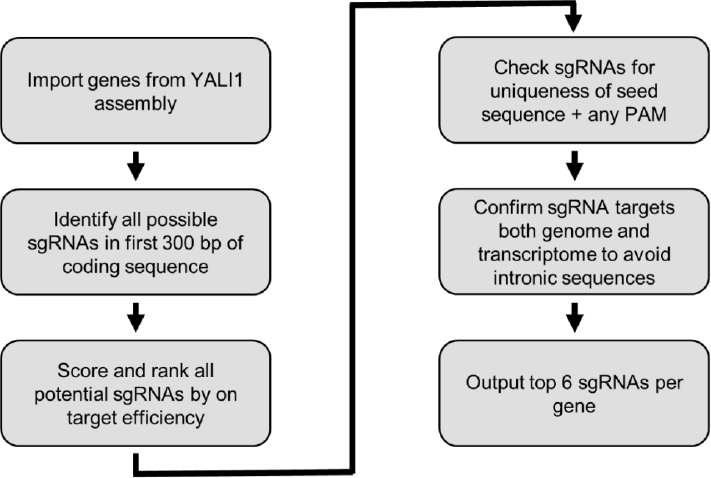
sgRNA design. Flowchart outlining sgRNA design strategy. On target efficiency calculated using previously constructed algorithm^1^. YALI1 is a recent assembly of the *Y. lipolytica* PO1f parent strain genome, W29^2^.

**Supplementary Figure 2.**
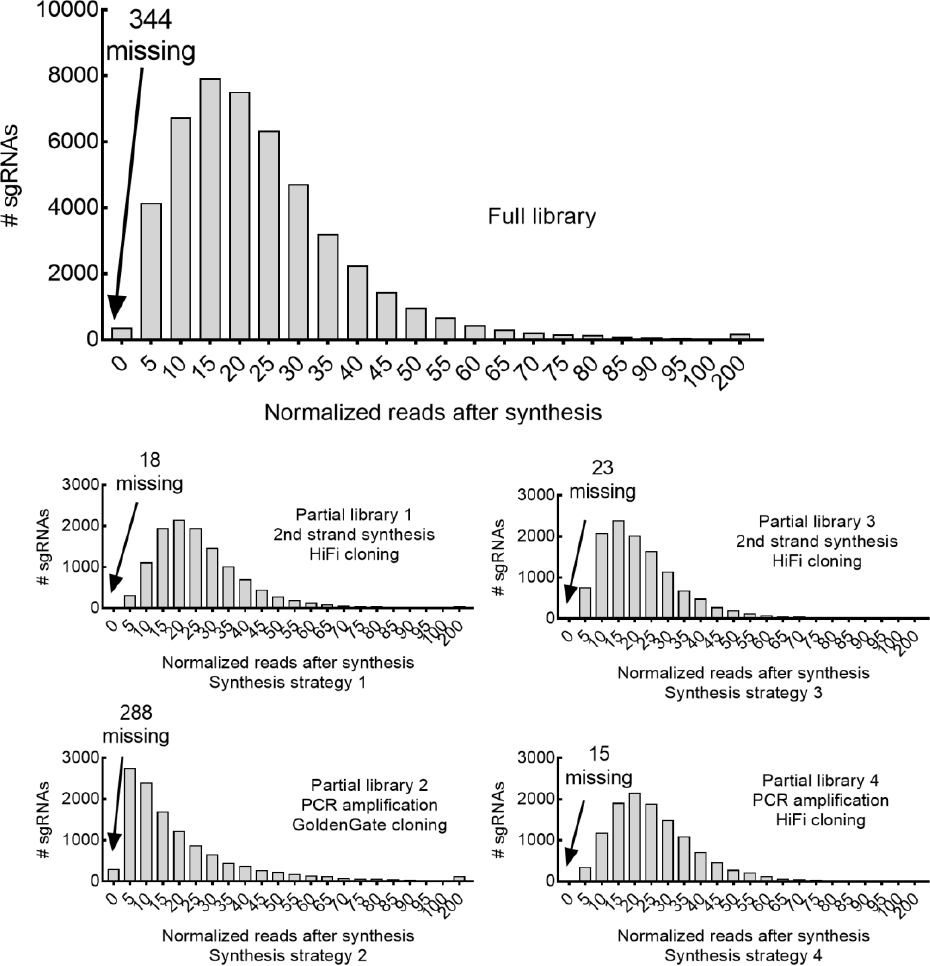
Library cloning results. sgRNA abundance distribution in the full library and for each partial library, which were generated using 3 different cloning strategies

**Supplementary Figure 3.**
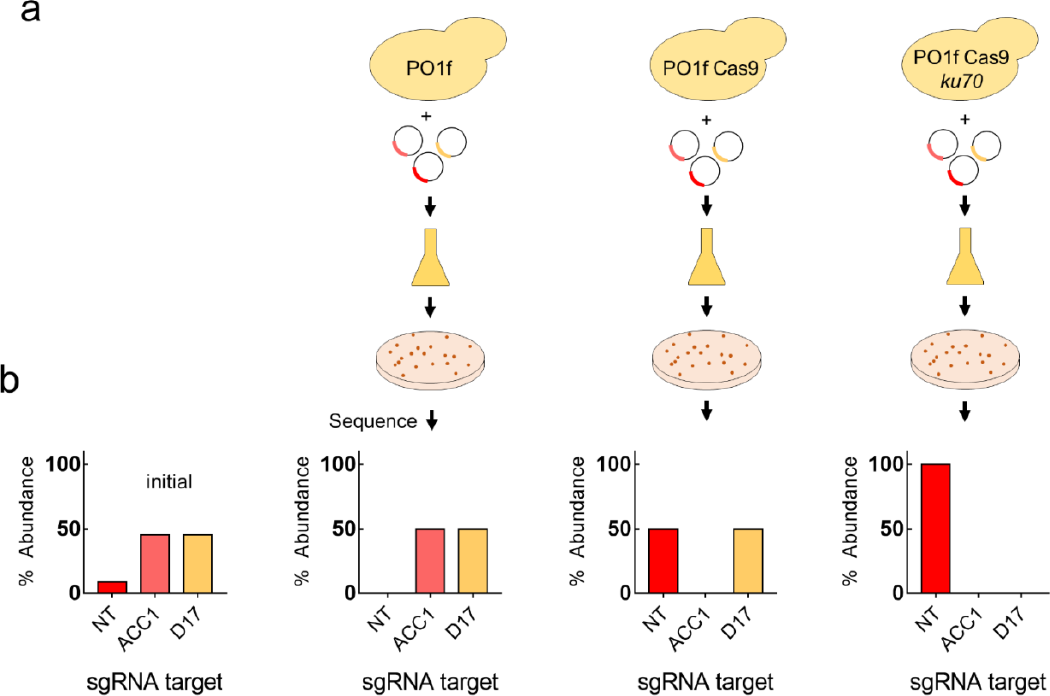
Pilot scale cutting and fitness score experiments. (a) Small scale transformation of PO1f, PO1f Cas9, and PO1f Cas9 *ku70* with 3 different sgRNA expression plasmids. *ACC1* (light red) is an essential gene, *D17* (yellow) is a validated nonessential gene, and *NT* (red) represents a nontargeting sgRNA. (b) Percent abundance of sgRNAs before (initial) and after transformation into each strain.

**Supplementary Figure 4.**
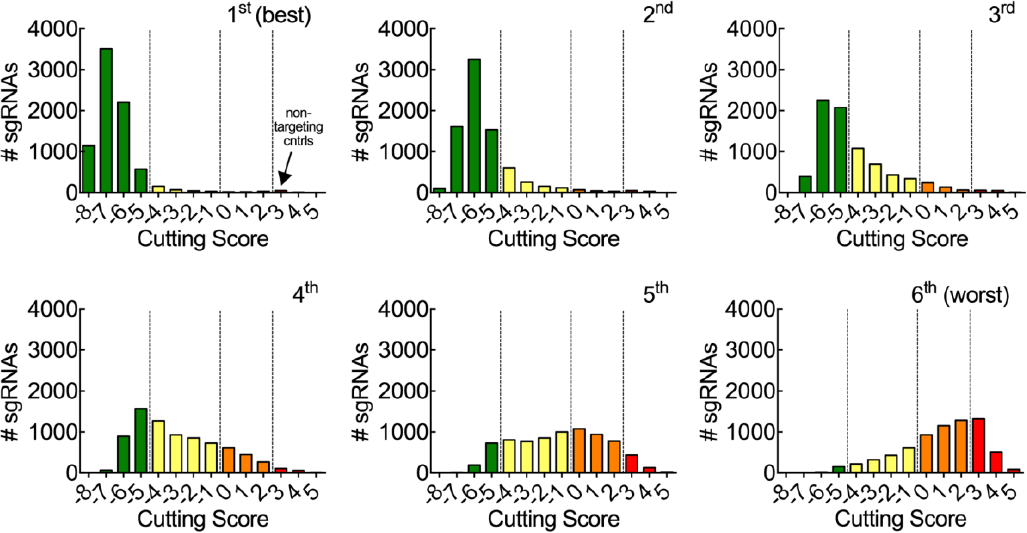
Distribution of cutting scores across library. The sgRNAs for each gene were ranked by CS from 1st to 6th and separated into six subpopulations each representing 1-fold genome coverage and assigned to sgRNA classes (Excellent (CS<−5; green;), Moderate (−5<CS<−1; yellow), Poor (−1<CS< 2; orange), and Noncutter (>2; red)). Mean CS values were determined after two growth cycles (4 days) from triplicate biological samples.

**Supplementary Figure 5.**
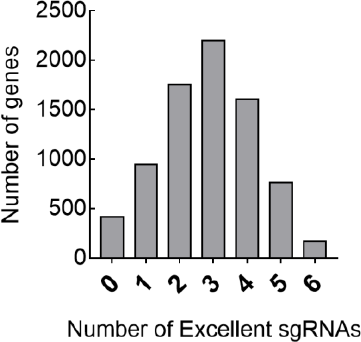
Excellent sgRNAs per gene. Frequency of the number of excellent sgRNAs for each gene in the genome.

**Supplementary Figure 6.**
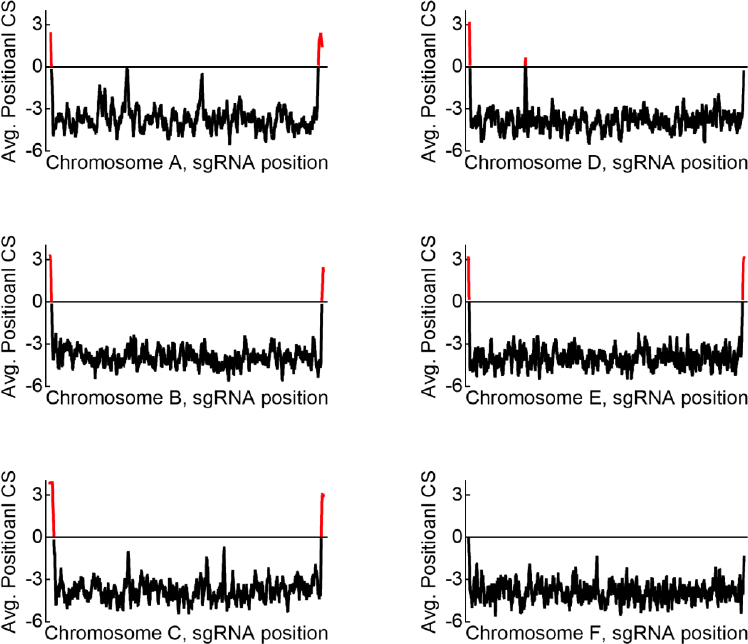
Cutting score by chromosome position. Moving average of cutting score across chromosomes. Average CS is calculated by moving average of 50 sgRNAs, ordered by position in each chromosome.

**Supplementary Figure 7.**
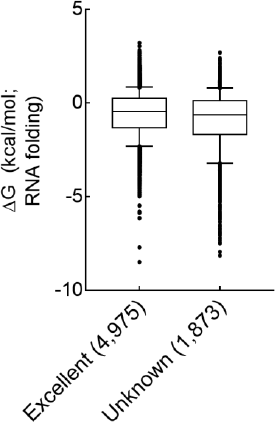
Secondary structure of excellent and noncutter sgRNAs. Predicted secondary structure^3^ for a representative population of excellent cutters and all noncutters that are not at chromosome ends or that contain a polyT motif.

**Supplementary Figure 8.**
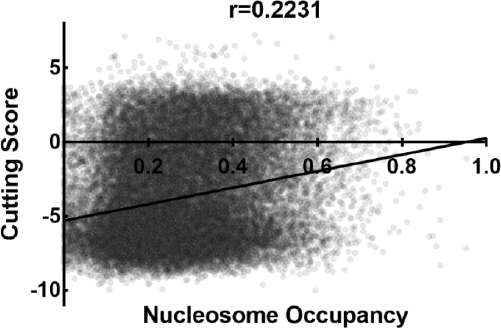
Comparison of nucleosome occupancy to cutting score. Plot of cutting score for each sgRNA compared to average nucleosome occupancy.

**Supplementary Figure 9.**
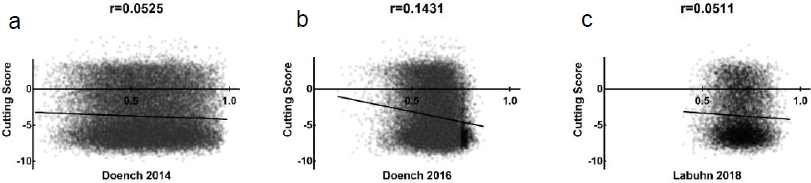
Comparison of sgRNA design algorithm score to cutting score. (a) sgRNA activity prediction from Doench *et al*. 2014^1^ compared to cutting score. (b) sgRNA activity prediction from Doench et al. 2016^4^ compared to cutting score. (c) sgRNA activity prediction from Labuhn et al. 2018^5^ compared to cutting score.

**Supplementary Figure 10.**
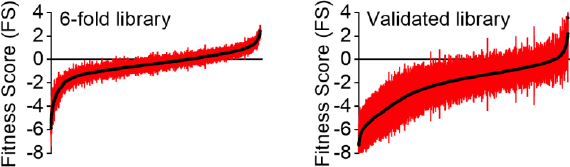
Rank-ordered curves of fitness scores with error. The mean and standard deviation of three biological replicates is shown for the full 6-fold coverage library and for the validated library.

**Supplementary Figure 11.**
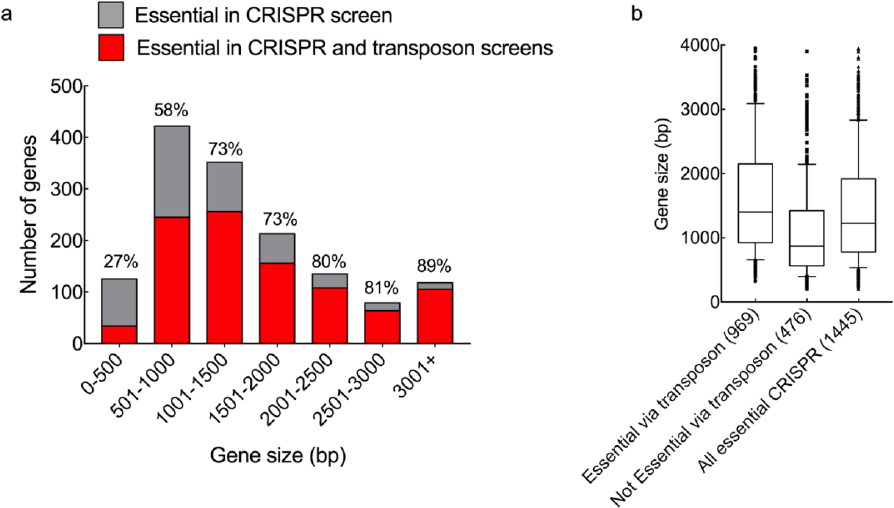
Transposon and CRISPR essential gene comparison. (a) Comparison of essential genes identified in CRISPR and transposon screens. All CRISPR essential genes are shown, with the percentages representing the amount that were also identified as essential in the transposon screen. (b) Size distribution of CRISPR essential genes that were also identified as essential by the transposon screen and those not identified as essential in the transposon screen.

**Supplementary Figure 12.**
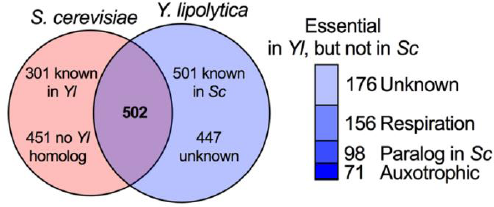
Essential gene comparison to *Saccharomyces cerevisiae*. Comparison of essential genes identified in *Y. lipolytica* with known essential genes from *S. cerevisiae*. Respiration indicates genes annotated as being essential for respirative growth in *S. cerevisiae*, Paralog in Sc indicates genes that are duplicated in the *S. cerevisiae* genome, and Auxotroph indicates genes that are involved in nucleotide or amino acid biosynthesis (identified via manual inspection). Unknown or no *Yl* homolog indicate genes that did not have a clear homolog.

**Supplementary Figure 13.**
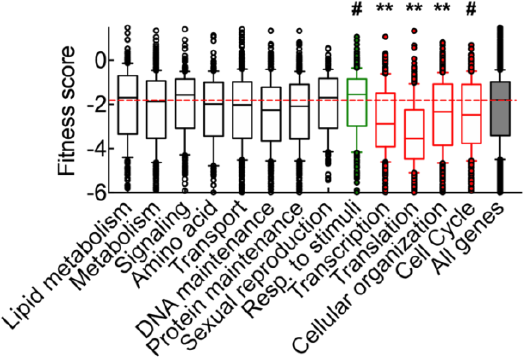
GO term analysis and fitness score. Fitness score of groups of genes separated by gene ontology classification of genes by *S. cerevisiae* homolog. Significance was determined by 1-way ANOVA with post hoc analysis (# p<0.05; * p<0.01; **p<0.0001).

**Supplementary Figure 14.**
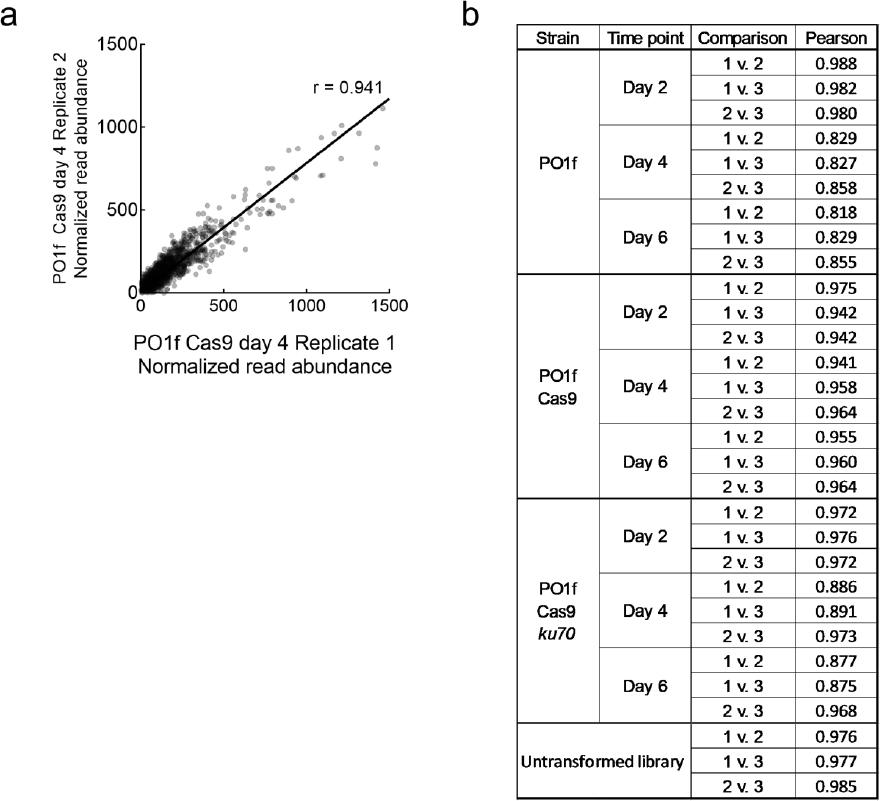
Pairwise comparison across biological replicates. (a) Pairwise comparison of PO1f Cas9 biological replicates 1 and 2. Linear regression analysis yielded a ‘Pearson’s coefficient of 0.941. (b) All possible pairwise comparisons of different replicates.

**Supplementary Table 1:**
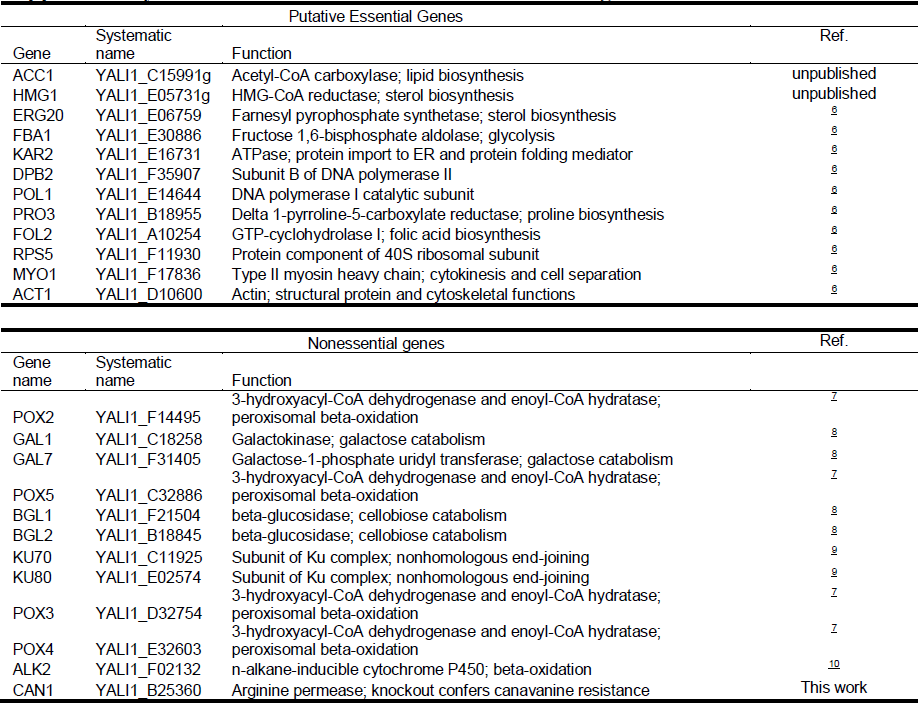
Selected essential and nonessential genes.

**Supplementary Table 2:**
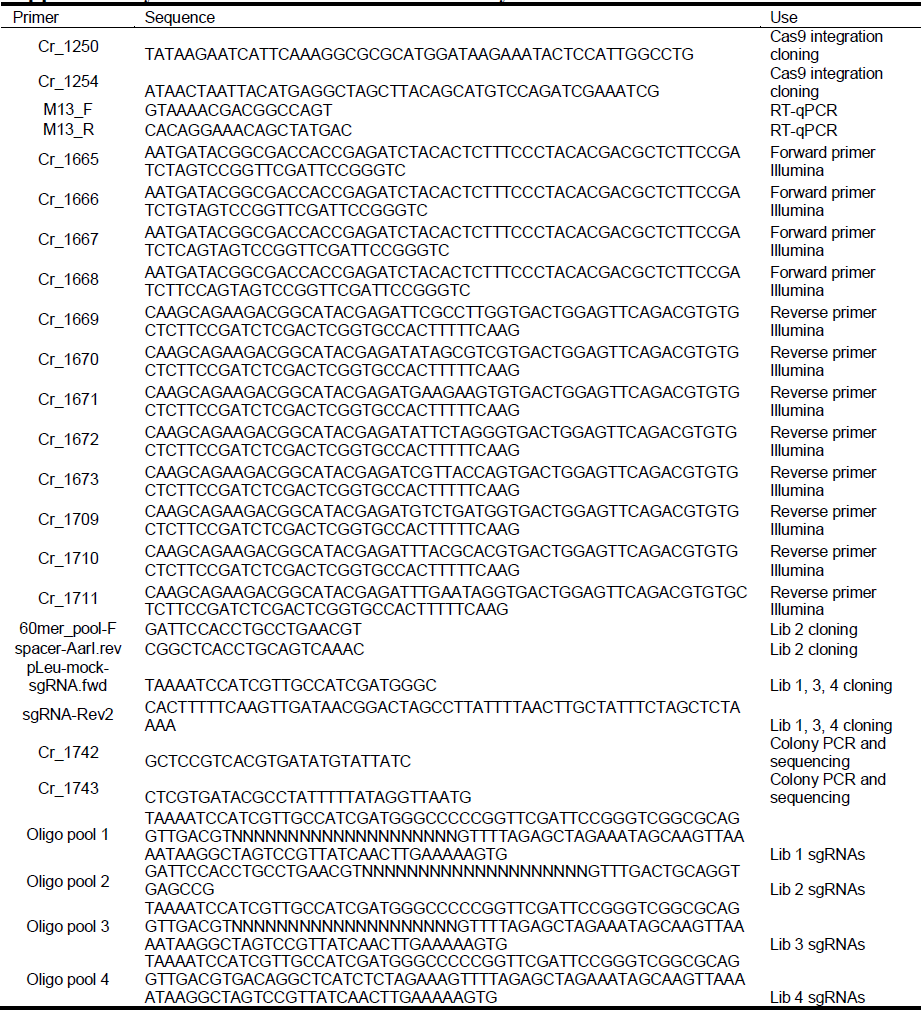
Primers used in this study.

